# Ultrafast immunostaining of organ-scale tissues for scalable proteomic phenotyping

**DOI:** 10.1101/660373

**Authors:** Dae Hee Yun, Young-Gyun Park, Jae Hun Cho, Lee Kamentsky, Nicholas B. Evans, Alex Albanese, Katherine Xie, Justin Swaney, Chang Ho Sohn, Yuxuan Tian, Qiangge Zhang, Gabi Drummond, Webster Guan, Nicholas DiNapoli, Heejin Choi, Hae-Yoon Jung, Luzdary Ruelas, Guoping Feng, Kwanghun Chung

## Abstract

Studying the function and dysfunction of complex biological systems necessitates comprehensive understanding of individual cells. Advancements in three-dimensional (3D) tissue processing and imaging modalities have enabled rapid visualization and phenotyping of cells in their spatial context. However, system-wide interrogation of individual cells within large intact tissue remains challenging, low throughput, and error-prone owing to the lack of robust labeling technologies. Here we introduce a rapid, versatile, and scalable method, eFLASH, that enables complete and uniform labeling of organ-scale tissue within one day. eFLASH dynamically modulates chemical transport and reaction kinetics to establish system-wide uniform labeling conditions throughout the day-long labeling period. This unique approach enables the same protocol to be compatible with a wide range of tissue types and probes, enabling combinatorial molecular phenotyping across different organs and species. We applied eFLASH to generate quantitative maps of various cell types in mouse brains. We also demonstrated multidimensional cell profiling in a marmoset brain block. We envision that eFLASH will spur holistic phenotyping of emerging animal models and disease models to help assess their functions and dysfunctions.

---

System-wide analysis of cell types is essential for understanding how complex cellular interactions give rise to various functions. Extensive efforts have been made towards characterizing cells, particularly in the brain, through various lenses (e.g., genomics, transcriptomics, proteomics, connectomics) and have established invaluable databases with new insights^1–7^. Among these approaches, proteomic imaging has distinct advantages. Mapping spatial distribution of proteins, the major functional substrate with distinct subcellular localization at single molecule precision, can provide rich molecular, functional, as well as morphological details of cells. Furthermore, visualizing endogenous proteins with highly specific antibodies does not require genetic manipulation or invasive *in vivo* surgery, and thus it is applicable to any species or tissue type including non-human primates and human clinical samples^8^.

When combined with intact organ transformation and clearing techniques, proteomic phenotyping can provide multiscale information, ranging from brain-wide cell distribution patterns to molecular and morphological details of individual cells without information loss caused by subsampling or 2D analysis^9–11^. However, scaling immunolabeling to large-scale tissues and higher species remains a major challenge in biology. Passive transport of large macromolecules such as antibodies into intact tissues can take weeks to months^9,12^. Antibody penetration can be further delayed or even blocked by target proteins with high expression levels, causing probe depletion and incomplete staining. Using excessive amounts of antibodies can improve probe penetration, but it becomes prohibitively expensive and thus unscalable. In conventional passive labeling approaches, experimental parameters for labeling (e.g., incubation time, probe amount) are highly dependent on sample properties (e.g., tissue type, size, shape) and target protein properties (e.g., expression level, distribution patterns), which are widely different between applications. Therefore, each experiment requires laborious, costly, and time-consuming optimization. The outcome of passive labeling is in many cases highly uneven with saturated labeling of outer regions and weak or no labeling of the core. Such non-uniform and incomplete labeling can prohibit automated analysis and cause systematic error. These challenges together have limited the power of 3D proteomic phenotyping to small tissues or a small number of established applications.

Here we present an integrated pipeline for holistic, rapid, scalable proteomic phenotyping of intact organs. To establish the pipeline, we developed an ultrafast and versatile immunolabeling technology, termed eFLASH (electrophoretically driven Fast Labeling using Affinity Sweeping in Hydrogel), which enables complete and uniform labeling of various types of tissues (mouse brain and intestine, human iPSC-derived cerebral organoid, and marmoset brain block) using a wide selection of antibodies (targeting structural, molecular, and neuronal activity markers) with a universal 1-day protocol. Combined with intact tissue processing and analysis techniques, we performed organ-wide quantification of various proteins at cellular resolution in mouse brains. We further demonstrated the power of 3D protein-based cell phenotyping by characterizing neural sub-types based on their 3D location, protein expression level, cell body size, and dendritic morphology in a fully integrated manner.

## RESULTS

### eFLASH mechanism

Our organ-wide molecular phenotyping framework consists of four major components: (1) intact tissue preservation via SHIELD, (2) volumetric labeling with eFLASH, (3) light-sheet imaging, and (4) automated 3D image analysis (Fig. 1a). The pipeline begins with robust preservation of biological tissue with SHIELD, which is a polyepoxide-based tissue fixation method that protects biomolecules and tissue architecture^13^. After rapid delipidation of the SHIELD-preserved tissues using stochastic electrotransport (SE)^12^, we immunolabel the intact tissues using eFLASH within just one day. The labeled samples are rapidly imaged at high-resolution using an axially swept light-sheet microscope. Finally, we analyze the resulting volumetric datasets via a suite of automated 3D image analysis algorithms to map various cell types within the tissue volume. Altogether, the pipeline enables extraction of organ-scale, single-cell-resolution data from a fresh sample within just 12 days (Fig. 1a).

**Figure 1.**
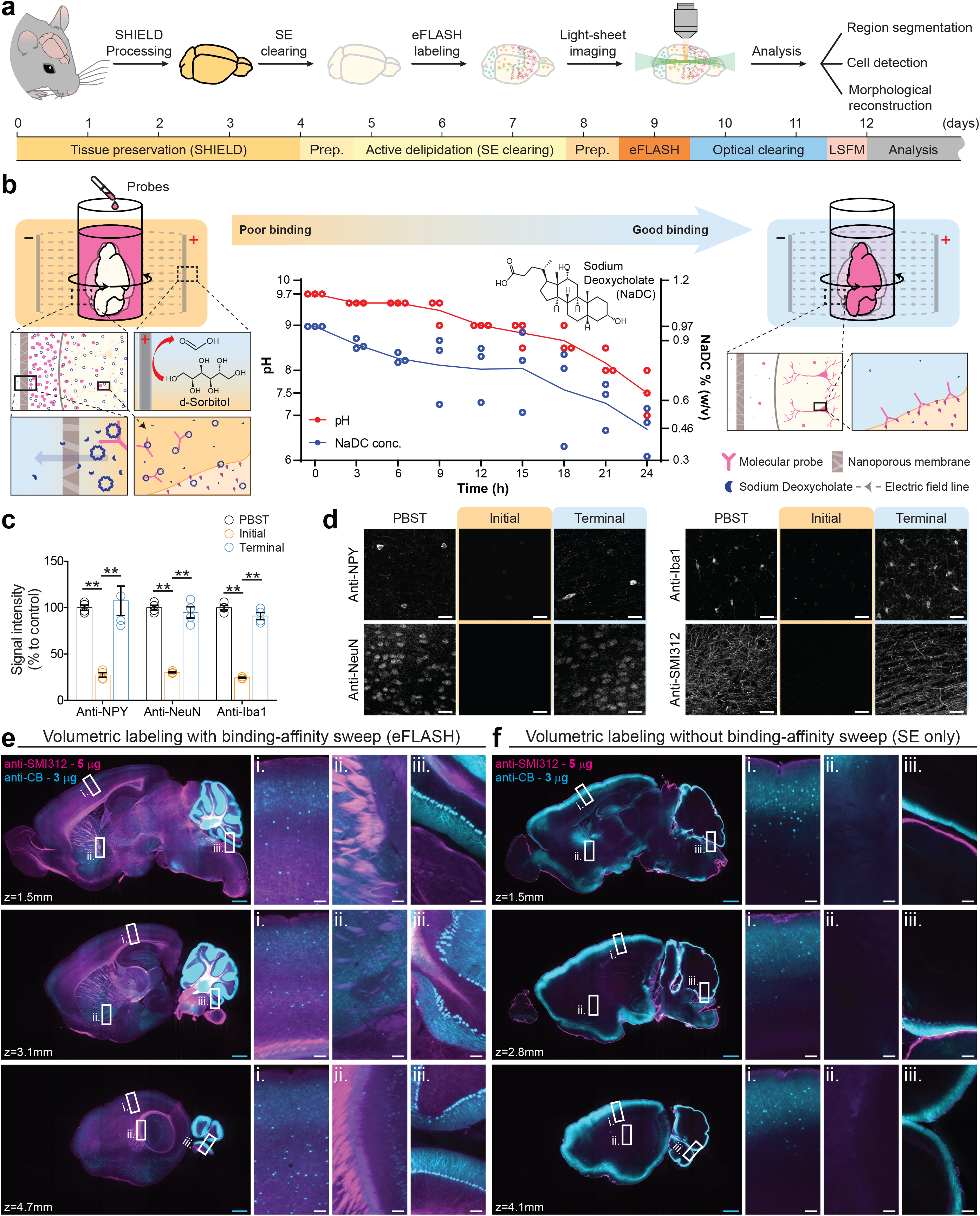
eFLASH enables rapid, uniform, and cost-efficient labeling of organ-scale tissues. (**a**) High-throughput pipeline for organ-wide molecular mapping at single cell resolution. The pipeline can generate high-resolution and multidimensional data from mouse brains within 12 days. SE, stochastic electrotransport; Prep, preparation; LSFM, light-sheet fluorescence microscopy. (**b**) eFLASH. The pH and sodium deoxycholate (NaDC) concentration of the labeling solution are gradually reduced to sweep the molecular probes’ binding affinity from unfavorable to favorable in the context of SE. Electrocatalytic oxidation of d-sorbitol on the anode surface generates acidic components that lower pH. NaDC concentration of the labeling solution is reduced by the concentration gradient through the nanoporous membrane. *N* = 3 independent experiments. Individual data points and mean. (**c**) Comparison of immunostaining signal among PBST control, initial (unfavorable binding condition) and terminal (favorable binding condition) eFLASH buffers. *N* = 4 tissue samples. Mean ± s.e.m.. Unpaired T-test, \*\**P* < 0.01. Scale bar = 50 μm. (**d**) Representative images used in (c). (**e-f**) Comparison of antibody penetration and uniformity of staining between eFLASH and SE only. Optical sections of mouse hemispheres at different depths are shown. The same amounts of antibodies were used for both experiments. Z-depth indicates the distance of optical sections from the mid-sagittal planes. Display ranges of images are 200-5,000 (cyan) and 100-500 (magenta) except **e-i**: 200-3,000 (cyan); **e-iii**: 200/10,000 (cyan) and 100/1,000 (magenta); **f-i**: 200/10,000 (cyan); **f-iii**: 200/20,000 (cyan) and 100/1,000 (magenta). Scale bars = 1 mm (cyan) or 100 μm (white).

eFLASH allows uniform immunolabeling of organ-scale tissues within a day by gradually shifting probe-target binding conditions from unfavorable to favorable while accelerating probe penetration using stochastic electrotransport (Fig. 1b). We discovered that bile salts, such as sodium deoxycholate (NaDC), can be used to control the labeling affinity for various antibodies in a concentration and pH dependent manner (Fig. 1c-d; Supplementary Fig. 1). A wide range of probes showed weak binding at high concentrations of NaDC and basic pH, but strong binding at low concentration of NaDC and neutral pH. These results indicate that labeling conditions can be gradually shifted from unfavorable to favorable by simultaneously sweeping pH (basic to neutral) and NaDC concentration (high to low).

To achieve a gradual pH sweep in an automated manner, we took advantage of electrochemical reactions that naturally occur under SE. Electrocatalytic oxidation of D-sorbitol produces acidic byproducts such as formic acid^14^. By adding D-sorbitol to a pH 9.5 buffer and letting it decompose by electro-oxidation under SE, we were able to gradually sweep pH from 9.5 to 7.5 over the course of one day (Fig. 1b, Supplementary Fig. 2).

Concentration of NaDC within the labeling solution was also swept in an automated manner using the concentration gradient established across a nanoporous membrane (Fig. 1b). The membrane, which separates the labeling solution and the outer solution, ensures that both molecular probes and large NaDC micelles remain within the labeling solution; however, it is permeable to NaDC monomers, small NaDC aggregates, and the rest of the buffer components. The initial concentration of 1% (w/v) NaDC within the labeling solution slowly decreases as the monomers travel down the concentration gradient to the outer solution, which contains 0.2% (w/v) of NaDC (Fig. 1b). We confirmed that the terminal buffer after pH and NaDC concentration shift allows strong antibody staining (Fig. 1c-d).

This progressive change in binding condition enables the probes to first penetrate deep into the tissue without being depleted and then increasingly bind to targets globally as the buffer composition gradually changes. This approach ensures uniform labeling of entire volumes regardless of the density and distribution pattern of the targets, specific binding kinetics of various antibodies, and the amount of antibody used. With eFLASH, even when using a minute amount of antibody for labeling highly dense targets (3 μg of antibody for calbindin and 5 μg of antibody for pan-axonal marker SMI312), high-quality uniform labeling could be achieved in a mouse brain hemisphere (Fig. 1e); however, without affinity-sweep, the small amount of antibody was quickly depleted on the surface before the core of the tissue could be labeled despite the increased transport speed provided by SE (Fig. 1f, Supplementary Video 1). These results indicate that eFLASH enables rapid, complete, and uniform immunolabeling of organ-scale tissues without the use of excessive amounts of molecular probes.

### Universal applicability of eFLASH

The affinity sweeping mechanism in eFLASH renders the technique insensitive to tissue type, size, or geometry. eFLASH is also insensitive to probe types because the sweeping range is wide enough to modulate binding affinities of many antibodies and other commonly used molecular probes. Therefore, the same operational parameters of eFLASH can be used for many applications without laborious and costly optimization. We found that a single protocol with the same parameters (e.g., voltage, pH, running time, chemical concentrations) is capable of uniformly labeling cerebral organoid, mouse intestine, mouse brain hemisphere, as well as marmoset brain block with various combinations of antibodies, allowing visualization of multiple proteins within a single sample (Fig 2a-d, Supplementary video 2-3).

**Figure 2.**
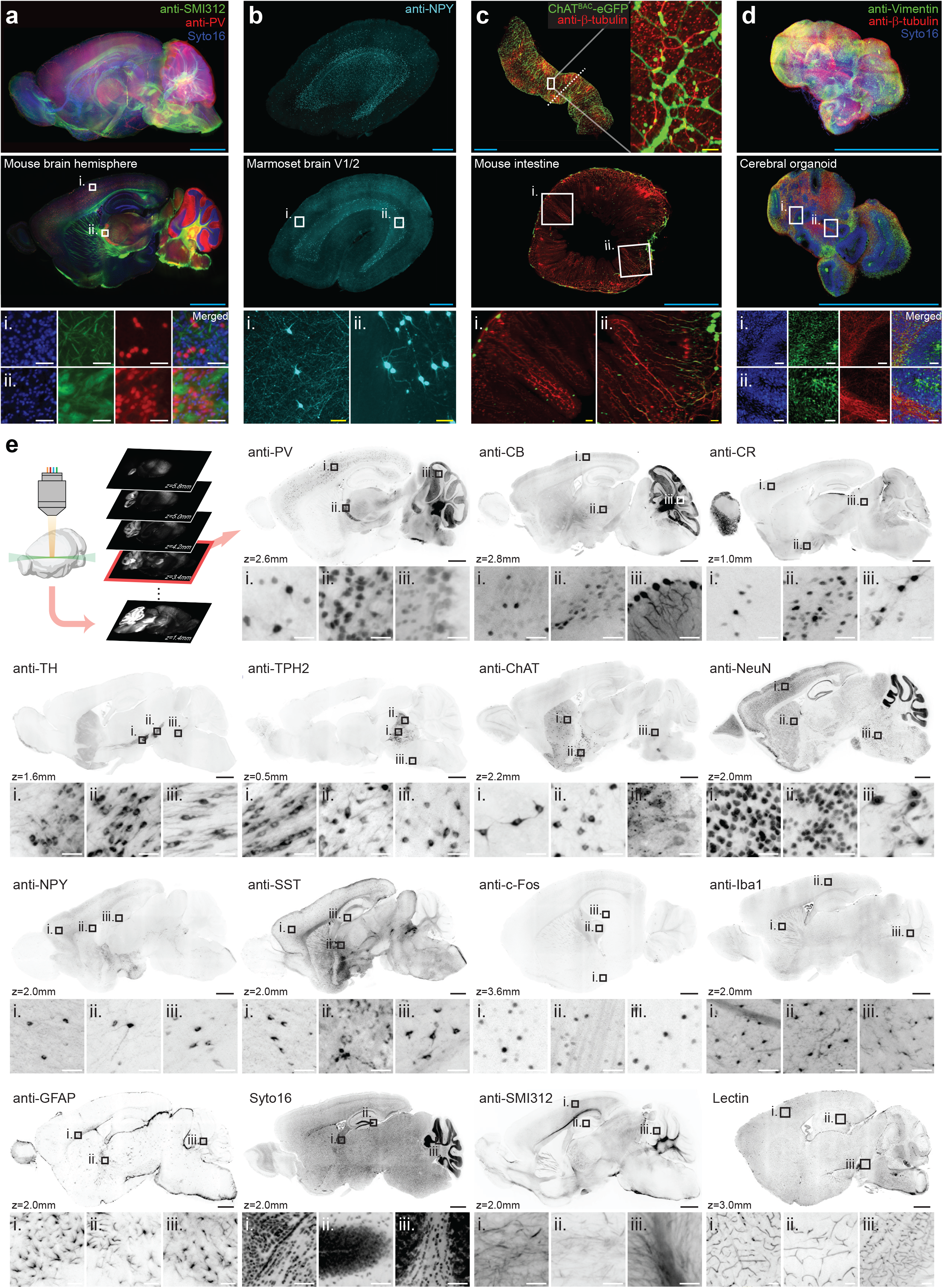
Single eFLASH protocol enables complete and uniform staining of various tissue types with a wide range of molecular probes. (**a-d**) Different types of tissue labeled with eFLASH. Full volume renderings (top row) and optical sections (middle and bottom row). Scale bars = 2 mm (cyan), 200 μm (yellow), and 50 μm (white). (**a**) An adult mouse brain hemisphere labeled with syto16 (blue), anti-SMI312 antibody (green), and anti-PV antibody (red). (**b**) A marmoset visual cortical block labeled with anti-NPY antibody (cyan). (**c**) ChAT^BAC^-eGFP mouse intestine labeled with anti-β-tubulin (red). (**d**) Cerebral organoid labeled with syto16 (blue), anti-Vimentin antibody (green), and anti-β-tubulin antibody (red). (**e**) Optical sections from whole adult mouse hemispheres labeled with indicated antibodies or molecular probes. Z-depth indicates the distance of the optical sections from mid-sagittal planes. PV, parvalbumin; CB, calbindin; CR, calretinin; TH, tyrosine hydroxylase; TPH2, tryptophan hydroxylase 2; ChAT, choline acetyltransferase; NeuN, neuronal nuclear antigen; NPY, neuropeptide Y; SST, somatostatin; Iba1, ionized calcium binding adaptor molecule 1. Scale bars = 1 mm (black) and 50 μm (white).

The same 1-day protocol was compatible with a wide range of antibodies harboring different binding affinities and target densities (Fig. 2e). eFLASH successfully labeled targets for various cell types (PV, CB, CR, NPY, SST, TH, TPH2, ChAT, NeuN, GFAP, Iba1), neuronal structure (SMI-312), and a neuronal activity marker (cFos) in intact mouse hemispheres (Fig. 2e, Supplementary video 4-5). The same eFLASH protocol was also compatible with lectin and Syto16, which are chemical probes that label blood vessels and nuclei, respectively. Together, these results suggest that eFLASH is a universal platform compatible with a wide range of tissue-types and molecular probes.

### A quantitative, brain-wide cell type mapping with eFLASH

eFLASH, combined with light-sheet microscopy, enables true volumetric quantification of protein expression at cellular resolution. eFLASH-stained mouse brain hemispheres were rapidly imaged using an axially swept light-sheet microscope at near-isometric resolution of 1.8 μm x 1.8 μm x 2 μm within 45 minutes per channel. Because the sample was processed and imaged as a whole without sectioning, the resulting volumetric data is an exhaustive representation of the sample that does not suffer from sampling errors and does not require interpolation or extrapolation to acquire brain-wide or region-specific cell counts. In addition, the multiplexed labeling capability of eFLASH allows analysis of cells co-expressing multiple proteins of interest with relative ease and flexibility compared to genetic labeling approaches. Currently, labeling up to four distinct targets is possible through transgenic labeling approaches^15^; however, developing transgenic mouse lines for each new combination of targets can be time consuming^16^.

To demonstrate the value of holistic labeling with eFLASH, we established an image analysis pipeline for atlas alignment, brain region segmentation, and cell detection for generating a quantitative map of various proteins. Volumetric images were automatically aligned to an atlas^4^ by linear and non-linear transformations based on Elastix^17^ then manually refined^18^. Each aligned 3D image volume was indexed to approximately 580 brain regions with 7 hierarchies. Brain-wide quantification of immunolabeled cells was accomplished using machine learning algorithms that were trained to identify features of individual cell-types (Supplementary Fig. 3). Specifically, Random Forest^19^ was applied after blob detection and principal component analysis (PCA). Detection performance was validated against manual ground-truth annotations of relevant brain regions that are known to express each cell type. Our cell detection pipeline achieved an f-score of higher than 90% for cortical regions and over 80% for subcortical brain regions for all tested cell-type markers. Using this pipeline, we were able to construct quantitative mouse brain atlases for multiple cell type defining makers, including CR, NPY, SST, TH, TPH2, and PV (Fig. 3a-c, Supplementary video 6).

**Figure 3.**
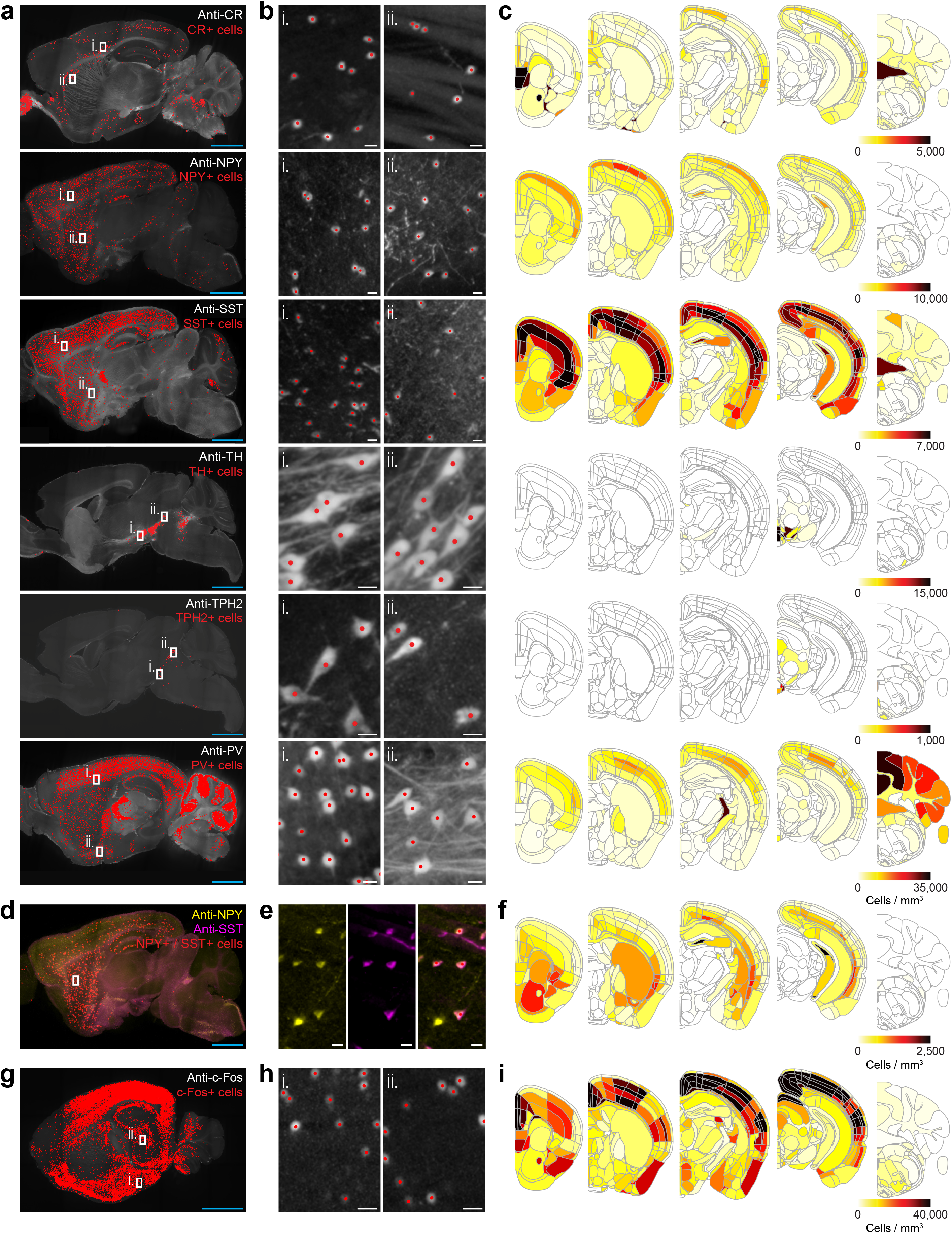
Quantitative brain-wide cell type mapping. (**a**) Optical section images of whole mouse hemisphere datasets. Adult mouse hemispheres were eFLASH-labeled with indicated antibodies and imaged. Automatically detected cells are marked with red dots. (**b**) Zoom-in views of **a.** (**c**) Representative images of 3D brain-wide cell type density heatmaps. See Supplementary video 6 for PV detection and heat map. (**d**) An optical section image of a whole mouse hemisphere co-labeled with anti-NPY antibody (yellow) and anti-SST antibody (magenta). NPY and SST co-positive cells are marked with red dots. (**e**) Zoom-in views of **d**. (**f**) Representative images of the 3D heatmap of the co-positive cells. (**g**) An optical section of a whole mouse hemisphere labeled with anti-c-Fos antibody. The mouse experienced contextual fear conditioning 90 minutes before sacrifice. (**h**) Zoom-in views of **g.** (**i**) Representative images of the 3D heatmap of c-Fos+ cells. Scale bars = 2 mm (cyan) and 20 μm (white).

The probe-insensitive nature of eFLASH enables co-labeling of multiple cell-types with any combinations. We performed simultaneous labeling of neuropeptide Y and somatostatin which are known to be co-expressed in a subset of GABAergic interneurons^20–22^ and of Tyrosine Hydroxylase and Tryptophan Hydroxylase 2 which are cell-type-specific markers for dopaminergic and serotonergic neurons, respectively, that are not generally known to overlap. In the case of NPY and SST, we mapped NPY+/SST−, NPY−/SST+, and NPY+/SST+ cells (Fig. 3 d-f). We found the highest density of NPY+ cells at layer 2 or 3 of the cerebral cortices (Fig. 3c), whereas SST+ cells showed the highest density at layer 4 or 5 in majority of the cortices^23^ (Fig. 3c) Interestingly, the highest density of cells that were co-positive for NPY and SST was seen in layer 5 or 6 (Fig. 3f). In a brain-wide average, 16 ± 4 % of NPY and 7 ± 5% of SST-expressing cortical cells were identified NPY+ / SST+ co-positive. In the case of TH and TPH2, we checked every TH+ and TPH2+ cells detected throughout the brain hemisphere and found that no cells were positive for both markers.

Finally, in addition to labeling cell-type defining proteins, brain-wide labeling of Immediate Early Genes (IEGs) such as c-Fos has been demonstrated as a powerful proxy for measuring neuronal activation^24,25^. We stained the brain of a mouse that experienced contextual fear conditioning 90 minutes before sacrifice with anti-cFos antibody and mapped its distribution (Fig. 3g-i). Examination of the dataset showed robust anti-c-Fos signal in hippocampus and amygdala areas, which are known to show increased activity upon contextual fear conditioning^26^. Combined, these results suggest that eFLASH-mediated immunolabeling can facilitate brain-wide quantification of protein expression at a cellular level in a high throughput and flexible manner.

### Brain-wide comparison of genetic and protein-based cell type labeling

Expression of genetically encoded fluorescent proteins have revolutionized biological labeling and imaging^27^, and ongoing developments in transgenic methodology offer powerful ways to study organ-wide gene expression^28–30^. However, the level of fluorescent protein expression is linked to transcription activity rather than the level of expression of mRNA or proteins, requiring careful and nuanced interpretation of data^31,32^. Additionally, several studies have reported on the importance of post-transcriptional processes that can often cause the quantities of mRNA and proteins to correlate poorly^32^, emphasizing the need for protein expression analysis.

Discrepancy between transgenic labeling and immunohistochemical labeling is widely recognized^28^, and there is a constant concerted effort to improve upon existing transgenic mouse lines for common targets^33–35^. To compare genetic labeling and eFLASH-mediated cell-type phenotyping approaches, we utilized transgenic mouse lines with two widely used transgene approaches: Cre-LoxP and BAC transgene^28,36–38^. First, we eFLASH-stained a hemisphere of a PV-Cre::DIO-tdTomato double transgenic mouse with anti-PV antibody (Fig. 4a). We performed the brain-wide quantitative analysis on tdTomato and anti-PV signals, and revealed substantial discrepancies between two labeling approaches, where the level of mismatch was highly heterogeneous among brain regions (Fig. 4b-c, Supplementary video 7). For example, in contrast to faithful tdTomato labeling of PV+ neurons in primary motor and primary somatosensory cortices (88% and 85% of tdTomato+ cells were also PV+), a substantial portion of tdTomato cells showed undetectable amounts of PV protein in some of cortical (e.g., 56% and 75% in the case of piriform and lateral entorhinal cortex) and subcortical (45% in caudate putamen; 62% in nucleus accumbens) areas. Furthermore, we found PV+ populations were not covered by the genetic labeling. For example, 66% and 77% of PV+ cells do not express tdTomato in CPu and NAc, respectively (Fig. 4b-c).

**Figure 4.**
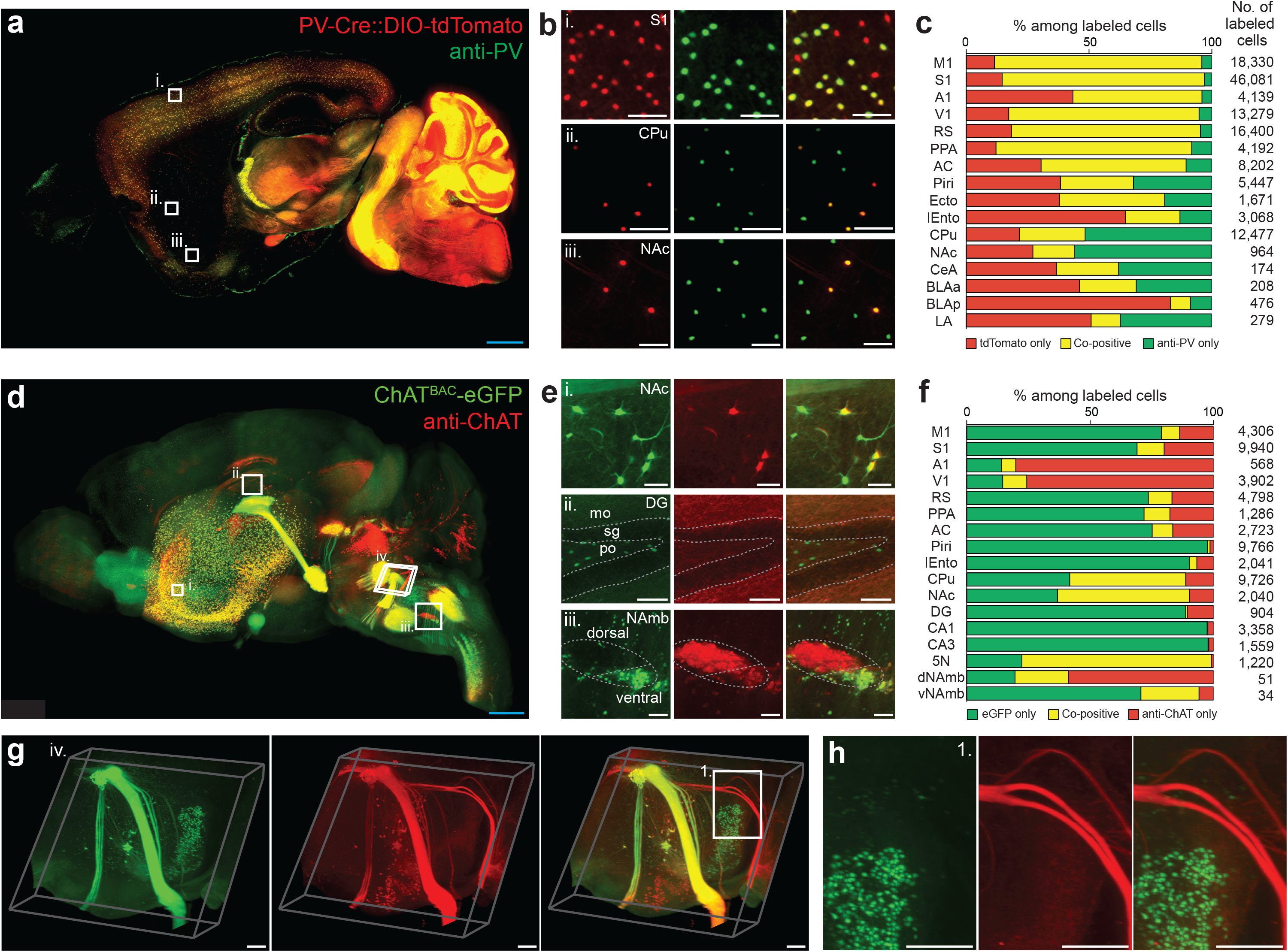
Brain-wide comparison of genetic cell-type labeling and eFLASH-driven protein-based cell type labeling. (**a**) An optical section of a 3D dataset from a PV-Cre and DIO-tdTomato dual transgenic mouse hemisphere stained with anti-PV antibody. (**b**) Zoom-in images of **a**. (**c**) A percentage plot for tdTomato-only (red), anti-PV-only (green), and tdTomato and anti-PV co-positive cells (yellow) among all the labeled cells in individual representative brain regions. (**d**) A 3D rendering of a ChAT^BAC^-eGFP mouse brain stained with anti-ChAT antibody. (**e**) Zoom-in views of **d**. (**f**) A percentage plot for eGFP-only (green), anti-ChAT-only (red), and eGFP and anti-ChAT co-positive cells (yellow) among all the labeled cells in individual representative brain regions. (**g**) Zoom-in view of **d**. (**h**) Zoom-in view of **g**. M1, primary motor cortex; S1, primary somatosensory cortex; A1, primary auditory cortex; V1, primary visual cortex; RSA, retrosplenial cortex; PPA, posterior parietal association cortex; AC, anterior cingulate cortex; Piri, piriform cortex; Ecto, ectorhinal cortex; lEnto, lateral entorhinal cortex; CPu, caudoputamen; NAc, nucleus accumbens; CeA, central amygdala; BLAa, basolateral amygdala, anterior part; BLAp, basolateral amygdala, posterior part; LA, lateral amygdala; DG, dentate gyrus; mo, dentate gyrus, molecular layer; sg, dentate gyrus, granule cell layer; po, dentate gyrus, polymorph layer; CA1, hippocampal CA1; CA3, hippocampal CA3; 5N, motor nucleus of trigeminal; dNAmb, nucleus ambiguus, dorsal part; vNAmb, nucleus ambiguus, ventral part. Scale bars = 1 mm (blue), 200 μm (white).

Next, we compared genetic and protein-based labeling of choline acetyltransferase (ChAT). eGFP expression via BAC transgene was highly divergent from the ChAT+ immunoreactivity pattern (Fig. 4d-h, Supplementary video 8). For example, only 9% and 14% of eGFP+ cells were also ChAT+ in M1 and S1 cortex. In the hippocampal CA1 and CA3, only 0.2% and 0.3% of eGFP+ cells showed detectable levels of ChAT immunoreactivity. Further, large populations of ChAT+ cells without eGFP expression were evident, especially in primary auditory and visual cortices (93%, 89%) (Fig. 4f). These discrepancies were heterogeneous even within the same brain region. Most ChAT+ cells were also eGFP+ in Nucleus ambiguus ventral part (80%), however, in its dorsal counterpart, only 26% of ChAT+ cells were colocalized with eGFP+ (Fig. 4e-iii). 3D visualization of the hemisphere also revealed labeling mismatch between fiber bundles. In the brain stem, we found a fiber bundle composed of ChAT+ axons without eGFP signals (Fig. 4g,h). These results suggest that eFLASH enables brain-wide analysis of transgenic labeling patterns and their validation by allowing simultaneous visualization of genetically expressed fluorescent proteins and immunolabeling signal within the same sample.

### Multidimensional single-cell analysis of marmoset visual cortex

Common marmoset *(Callithrix jacchus*), a small New World primate, has emerged as a powerful model for neuroscience research^39^. Their rapid reproduction cycles and compatibility with existing genetic engineering tools renders them a promising model for studying various brain disorders. Holistic cell-level phenotyping of the marmoset brain, however, remains challenging owing to the limited quality and availability of transgenic lines, significantly higher cost and larger brain size compared to rodent models.

Protein-based cellular phenotyping using eFLASH and SHIELD can be advantageous for higher model systems, including primates, where genetic manipulation remains challenging^40,41^. Moreover, the multiplexing capability of this approach allows simultaneous mapping of various molecular and cell-type markers within the same brain tissue, which not only increases the dimensionality of integrated phenotypic analysis, but also decreases the number of animals required for a study and consequently the cost.

To test this idea, we applied eFLASH and SHIELD to characterize cells in an intact marmoset brain block of visual cortex (5 mm x 5 mm x 8 mm). First, we eFLASH-stained the SHIELD-preserved sample with anti-PV antibody. From the holistic visualization and detection of PV+ cells in the sample (Fig. 5a), we found that the inter-layer distribution of PV+ cell is heterogeneous among parts of the visual cortex. The density of PV+ cells was higher in the area facing the calcarine sulcus compared to the other area of the visual cortical block (1770.3 ± 56.4 vs 979.0 ± 33.4 cells per mm^3^, unpaired T-test, P<0.0005, N= 4 of 120 μm-thick optical sections) (Fig. 5b-c, Supplementary video 9). We also observed that several cortical areas were devoid of PV+ neurons (Fig. 5a-c). Furthermore, we observed that inter-layer distribution patterns of PV+ cells differed between mouse and marmoset visual cortex (Fig. 5c). After mapping PV+ cells, we destained the same marmoset brain block and re-stained it with anti-NPY antibody using eFLASH. We found that NPY+ cells are mostly localized in layer 6 and white matter of the marmoset visual cortex, which was in contrast with mouse visual cortex that showed a more uniform NPY+ cell distribution across the cortical layers (Fig. 5d-f).

**Figure 5.**
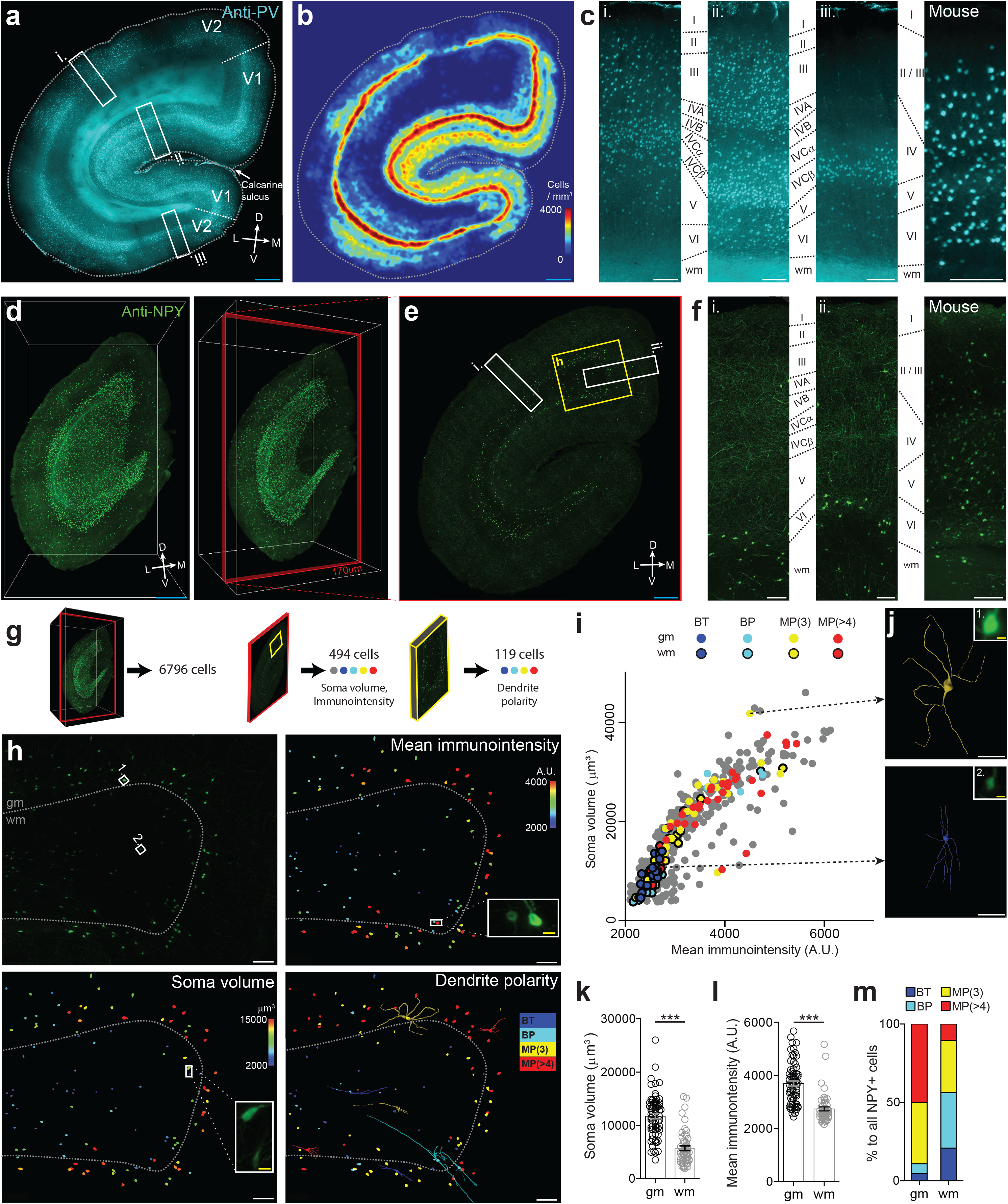
Multidimensional analysis of an eFLASH-stained marmoset brain block at single cell resolution. (**a**) A marmoset visual cortical tissue (5 mm x 5 mm x 8 mm) was eFLASH-stained with anti-PV antibody. An optical section of the 3D block image is shown. D, dorsal; V, ventral; L, lateral; M, medial. (**b**) PV+ cell density analysis shown as heat map. See Supplementary video 9 for volumetric heat map. (**c**) Inter-layer distribution of PV+ cell in marmoset (**i-iii**) and mouse visual cortex (right). wm, white matter. Zoomed regions indicated on panel **a**. (**d**) The same marmoset brain block was destained then re-stained with anti-NPY antibody with eFLASH. 3D volume renderings from two different perspectives. (**e**) A coronal optical section from the 3D data in **d.** (**f**) Inter-layer distribution of NPY+ cells in marmoset (**i-**ii) and mouse visual cortex (*right*). Zoomed regions indicated on panel **e**. (**g**) 6796 cells were detected in the 3D volume. Soma volume and mean immunointensity analysis was performed on 494 cells within the selected 170 μm-thick optical section (*red*). Dendrite polarity analysis was performed on 119 cells within the indicated volume (*yellow*). (**h**) An intensity projection image of a cortical fold region indicated on panel **e** (*upper left*). All cells within this volume were analyzed for their soma volume (*lower left*), mean immunointensity (*upper right*), and dendrite polarity with dendrite traces shown for 8 representative neurons (*lower right*). gm, gray matter; A.U., arbitrary unit; BT, bitufted cell; BP, bipolar cell; MP (3), multipolar cell with 3 primary dendrites; MP (>4), multipolar cell with 4 or more primary dendrites. (**i**) Annotated NPY+ cells plotted based on their mean immunointensity and soma volume. Total cells plotted, N = 494. (**j**) 3D reconstruction of dendrites of gray (*upper*) and white matter NPY+ cell (*lower*) in **h**. Insets indicate soma image of each cell. (**k**-**l**) Soma volume (**k**) and mean immunointensity (**l**) of gray and white matter NPY+ cells in **h**. N = 67 and 52 for gray and white matter cells, respectively. (**m**) Percentages of NPY+ cells categorized by their dendrite polarity. Mann-Whitney test, *P < 0.05, **P < 0.01, ***P < 0.005. Mean ± s.e.m.. Scale bars = 1 mm (blue), 200 μm (white), 20 μm (yellow).

Immunostaining can provide access to cellular morphology without genetic labeling or dye injections because many proteins are distributed or transported to cytoplasm and subcellular compartments. Using eFLASH-mediated volume-wide immunolabeling, we may be able to characterize both morphological and molecular details of individual cells throughout intact tissue volumes. To demonstrate this possibility, we performed deep analysis of individual NPY+ cells in a cortical fold sub-volume. From the automatically detected 6796 NPY+ cells in the volume, we quantified the soma volume and mean immunointensity of 494 cells, and dendrite polarity of 119 cells (Fig. 5g-i). Analysis of dendritic morphology of individual NPY+ cells led us to classify the cells into separate categories based on previously established descriptions of GABAergic interneurons: bitufted, bipolar or multipolar^42^ (Fig 5h, Supplementary Fig 4.) Compared to NPY+ cells in white matter, NPY+ cells in gray matter have soma with larger volume and higher mean fluorescent signal intensity (Fig. 5k-l), suggesting higher intracellular concentration of NPY protein^43,44^. We also found that most gray matter NPY+ cells are multipolar cells, whereas most NPY+ cells in white matter were bitufted or bipolar cells (Fig. 5h,j,m; Supplementary video 10). Together, these results demonstrate that complete and uniform immunolabeling of large-scale intact tissues with eFLASH enables high-dimensional phenotyping of individual cells even on model animals with limited access to genetic tools.

## DISCUSSION

In this study, we developed a rapid, versatile, and scalable immunolabeling technology, eFLASH, that enables complete and uniform immunolabeling of organ-scale tissues within one day for protein-based high dimensional cellular phenotyping. The universal 1-day protocol based on the gradual sweeping of probe-target binding affinity allows labeling of various markers simultaneously for disparate tissue types. Combined with the volumetric imaging and analysis pipeline, eFLASH enables 3D visualization and multi-dimensional phenotyping of molecular markers in large intact tissues with single-cell-resolution.

eFLASH is rationally designed to address the main challenge in scaling molecular labeling to organ-scale samples: the drastic mismatch between probe diffusion time scale and probe-target reaction time scale. Probe-target binding reaction is orders of magnitude faster than probe diffusion^12^. The diffusion timescale increases quadratically with the thickness of the sample, whereas probes rapidly bind to targets as soon as they encounter. If the density of the target molecule is high, which is the case for many of protein targets, probes cannot penetrate deeper into the tissue until they saturate all target molecules in their path. This means that uniform and complete labeling of intact tissue is not possible without using a large amount of probes, reducing the tissue size, or reducing the density of antigens.

Transport of electromobile molecules such as antibodies can be expedited using stochastic electrotransport^12^. However, the probe transport time scale in SE is still much longer than the reaction time scale. Applying a higher electric field can further increase transport speed, but Joule heating can cause tissue damage. Therefore, it is imperative to modulate both the rate of reaction and transport simultaneously. Switching off the binding reaction allows transport of antibodies into the core of the tissue without depletion^45^. Once probes reach the core of the sample, the binding reaction can be switched back on by changing the surrounding chemical environment (pH, detergent concentration). Discrete modulation of kinetics by such step-wise change, however, inevitably forms concentration gradients of chemicals (e.g., pH and NaDC) and probes inside the tissue, which causes uneven labeling. We addressed this challenge in eFLASH by slowly and gradually changing the concentration of the chemicals to ensure that the reaction condition is uniform tissue-wide throughout the day-long labeling period.

eFLASH is a robust process that offers considerable experimental flexibility. Repeated staining of the same tissue is possible with eFLASH, allowing multiple interrogations of precious samples as demonstrated with the marmoset brain block (Fig. 5). eFLASH can also be used to immunolabel the organs of transgenic mice expressing fluorescent proteins, allowing simultaneous visualization of both genetic labeling and immunolabeling signals (Fig. 2c, Fig. 4). This suggests that eFLASH can be utilized for comprehensive immunohistological validation of genetic labeling, amplification of genetically labeled signal using anti-fluorescent protein antibodies, and multiplexed proteomic analysis of genetically labeled cells in intact tissues.

Recently, tissue-clearing techniques and volume imaging methods have been applied to whole organ samples to demonstrate the potential of 3D phenotyping with single-cell resolution^24,46,47^. Many of these studies utilized genetic labeling which provides both uniform and high signal-to-noise ratio suitable for computational analysis^29^. However, genetic labeling is relatively inflexible when it comes to target selection, as new transgenic mouse or protocol is required for each target or each combination of targets^16,36^. With eFLASH, the choice of targets and the combinations of targets is based simply on the availability of compatible molecular probes. Additionally, eFLASH performs direct immunohistological labeling of target proteins present in the tissue, allowing for simplified interpretation of resulting data. As powerful as it is, Cre-LoxP transgenesis is known to suffer from false-positive (e.g., transgene-independent CRE expression, CRE-independent recombination) and false-negative labeling (e.g., CRE mosaicism)^48^. We observed that there was a discrepancy between fluorescent protein signal and antibody labeling signal in PV-Cre:DIO-tdTomato double-transgenic mouse brain labeled with anti-PV antibody (Fig. 4), Reversible tissue and temporal specific control systems, (such as a tetracycline response system), and BAC transgenesis resolved some of these issues, but not all. For example, discrepancies between genetic and protein-based labeling in BAC transgenic mouse lines were observed in both previous^28,34,49^ and present studies (Fig. 4), and it has been suggested that expression of BAC transgene can be affected by the presence of other transcription factors, microRNAs, or control regions of gene fragments^16,50^. We anticipate that protein-based mapping enabled by eFLASH can complement the cutting-edge genetic labeling approaches (e.g., viral labeling) for anatomical, molecular, and functional mapping of neural circuits.

Furthermore, eFLASH can facilitate studies of animal models with limited access to genetic labeling methodologies. The Common Marmoset is an emerging primate model for social behaviors with many experimental advantages^51,52^, and thus much effort has been undertaken to construct marmoset brain atlases in diverse modalities^53–55^. Unfortunately, numerous hurdles remain in translating existing genetic labeling approaches for rodents to marmosets. For example, germline genetic manipulation for generating transgenic primates is still difficult and expensive^40,56^, and long gestation and maturation period of primates as well as ethical concerns make each primate sample highly precious. Viral labeling approaches have shown the most promise; however, clear limitations exist since most enhancer elements are not defined and viral vectors have limited capacity to include large gene elements^40^. Moreover, achieving systemic coverage of the entire brain with viral labeling also remains challenging^57^. We anticipate that the scalability and flexibility of eFLASH will aid organ-wide phenotyping efforts on such model animals.

We envision that the versatility and high throughput capabilities of eFLASH will benefit numerous studies requiring system-wide yet highly detailed views of biological tissues, especially for exploratory studies comparing healthy and diseased animals or of model animals with limited access to genetic labeling strategies. Application of eFLASH will synergize greatly with advancements in biological imaging, molecular binder technologies, and computational frameworks for big data analysis^58^. Holistic, rapid, and unbiased approaches enabled by such synergistic technological advances will ultimately aid in providing a broader perspective in the study of complex biological systems.

## Acknowledgements

The authors thank the entire Chung laboratory for support and discussions. We acknowledge N. Peat for contribution to chemical screening for buffer development. K.C. was supported by the Burroughs Wellcome Fund Career Awards at the Scientific Interface, Searle Scholars Program. Packard award in Science and Engineering, NARSAD Young Investigator Award, and the McKight Foundation Technology Award. This work was supported by the JPB Foundation (PIIF and PNDRF), the NCSOFT Cultural Foundation, the Institute for Basic Science IBS-R026-D1, and the NIH (1-DP2-ES027992, U01MH117072).

## Author Contributions

D.H.Y., Y.-G.P., J.H.C. and K.C. designed the experiments and wrote the paper with input from other authors. D.H.Y., J.H.C., and K.C. designed eFLASH protocols and systems. D.H.Y. and J.H.C. performed the volumetric labeling experiments with N.D.’s help. Y.-G.P. aided the development of the eFLASH technology by performing passive staining experiments for screening antibodies and buffers, and imaging eFLASH-labeled samples. Y.-G.P. led SHIELD-processing of all tissue samples with K.X.’s help. G.F. and K.C. initiated the marmoset brain mapping project. G.F. provided the marmoset and Q.Z. perfused the marmoset. L.K. and J.S. developed the computational pipeline with Y.-G.P., D.H.Y., W.G., and K.C.’s input. N.B.E. and Y.-G.P. performed light-sheet imaging with H.C.’s help. D.H.Y. performed active delipidation of mouse and marmoset samples with N.D.’s help. D.H.Y. performed the buffer characterization in Figure 1. A.A. provided and imaged the SHIELD processed cerebral organoid for Figure 2. Y.-G.P. and L.K. performed brain-wide cell-type mapping in Figure 3 with D.H.Y. and K.X.’s help. Y.-G.P. performed co-positivity analysis for Figure 4 and the multi-dimensional analysis of marmoset datasets in Figure 5. C.H.S. aided in antibody and fluorescent dye screening for the project. G.D. and Y.X. helped with initial manuscript preparation. Y.X., H.-Y.J., and L.R. aided in detergent and buffer screening and characterization.

## Online Methods

### Mice

Young adult (2-4 month) C57BL/6 mice were housed in a 12 hr light/dark cycle with unrestricted access to food and water. All experimental protocols were approved by the MIT Institutional Animal Care and Use Committee and the Division of Comparative Medicine and were in accordance with guidelines from the National Institute of Health. The following transgenic lines were used for this study: Thy1::GFP M-line, Thy1::YFP H-line, ChAT^BAC^-eGFP (Jackson Stock No. 007902), PV-Cre / loxP-tdTomato (Jackson Stock No. 017320 and 007914), and Fos-CreER^T2^/ DIO-tdTomato (Jackson Stock No. 021882, 007914).

### Marmoset

All animal experiments were approved by the Institutional Animal Care and Use Committee of Massachusetts Institute of Technology and were performed under the guidelines from the National Institute of Health. Adult common marmosets (2-4 years old) were housed in AAALAC-accredited facilities. The housing room was maintained at 74.0 ± 2.0 °F (23.3 ± 1.1 °C), in the relative humidity of 50 ± 20 %, and in a 12 hr light/dark cycle. The animals were housed in dedicated cages with enrichment devices and had unrestricted access to food and water.

For histological examinations, the animals were deeply sedated by intramuscular injection of Ketamine (20-40 mg/kg) or Alfaxalone (5-10 mg/kg), followed by intravenous injection of sodium pentobarbital (10-30 mg/kg). When pedal withdrawal reflex was eliminated and/or respiratory rate was diminished, animals were perfused transcardially with 0.5 ml 1000 IU/ml heparin and 100-200 ml cold PBS by gravity. Then the descending aorta of the animals was clamped, and a peristaltic pump was used to infuse another 200-300 ml ice-cold SHIELD perfusion solution (10%(w/v) GE38 and 4% PFA(w/v) in PBS). Brains were removed from the skulls and SHIELD-processed.

### Organoids

Organoids were grown according to the protocol by Lancaster et al.^59^, with the addition of dual SMAD inhibition between d6 and d9 to increase neural differentiation as previously described^60^. Organoids were grown from iPSC cells (System Biosciences, #SC101A-1). After Matrigel droplet embedding, organoids were transferred to 60 mm suspension culture dishes (Corning, #430589) and placed on shaker at 75 rpm on day 16. The organoids were SHIELD-processed at day 35 (see the section “SHIELD processing”).

### Contextual fear conditioning

Contextual fear conditioning (CFC) was conducted using a chamber with an animal shocker (Habitest, Coulbourn, MA). After 300 s exploration in the chamber, mice were shocked (0.75 mA, 2 s) and maintained in the chamber 5 minutes more. Mice were sacrificed 60 minutes after the behavioral test was ended.

### Sodium Deoxycholate (NaDC) concentration measurement

Concentration of surfactants can be measured by the degree of solubilization of hydrophobic organic dyes. Above the critical micelle concentration, the amount of solubilized dye increases linearly with the increase in surfactant concentration^61^. Degree of solubilization was measured based on light absorption using a microplate reader at 505nm. Sufficient Orange OT dye (Sigma, 344664, powder) was added to fully saturate 200-proof ethanol at RT. 200 μl of saturated solution was added to each of the wells in 96-well plate and allowed to fully evaporate to deposit Orange OT dye to the well surface. 100 μl of eFLASH buffer collected at various time points were added to the prepared wells and left on an orbital shaker overnight. The well plate was centrifuged at 2000g for 10 minutes (Multifuge X1R, Thermofisher). 50 μl from each well was collected and added to a black 96-well plate with glass bottom for measurement using a microplate reader (EnSpire Multimode Plate Reader, Perkinelmer). NaDC concentration was calculated based on a standard curve generated using the method described above from solutions with known concentrations of NaDC (Supplementary Figure 2).

### SHIELD processing

Preservation of mouse brain hemispheres were carried out according to the previously published SHIELD protocol^3^. Mice were transcardially perfused with ice-cold PBS and then with the SHIELD perfusion solution. Dissected brains or organs were incubated in the same perfusion solution at 4 °C for 48 h. Tissues were then transferred to the SHIELD-OFF solution (1X PBS containing 10% (w/v) P3PE) and incubated at 4 °C for 24 h. In the case of brain hemisphere processing, a whole brain was split into hemispheres before being incubated in the SHIELD-OFF solution. Following the SHIELD-OFF step, the organs were placed in the SHIELD-ON solution (0.1 M sodium carbonate buffer at pH 10) and incubated at 37 °C for 24 h.

Marmoset brains perfused with ice-cold PBS and then with SHIELD perfusion solution were incubated in the same perfusion solution at 4 °C for 48 h. The brain was hemisected, transferred to the SHIELD-OFF solution, and incubated at 4 °C for 24 h. Following the SHIELD-OFF step, the hemispheres were placed in the SHIELD-ON solution and incubated at 37 °C for 24 h. Afterwards the hemispheres were transferred to PBS for washing.

Organoids were fixed in 1X PBS with 4% (w/v) PFA at RT for 30 minutes and subsequently incubated in SHIELD-OFF solution at 4°C for 48h. Samples were then incubated in SHIELD-ON solution at 37°C overnight before washing with PBS with 0.02% sodium azide at RT for at least 24 h.

### Passive clearing (delipidation)

SHIELD-processed samples were delipidated before labeling or imaging. Passive delipidation was done by incubating tissues in the clearing buffer (300 mM SDS, 10 mM sodium borate, 100mM sodium sulfite, pH 9.0). Thin slices between 100 μm and 200 μm thickness were cleared at 45°C clearing buffer for 2-3 hrs. Mouse brain hemispheres were cleared at 45°C for 10-14 days. Organoids were cleared at 55°C for 36 hrs.

### Active clearing (Stochastic Electrotransport)

SHIELD-processed samples can also be cleared rapidly using stochastic electrotransport (SmartClear Pro, LifeCanvas Technologies). Mouse brain hemispheres were cleared at 45°C for 3-4 days. The marmoset brain hemisphere was cut coronally into 4 blocks of 8 mm-thickness using a microtome and the blocks were cleared at 45°C for 2 weeks.

### Antibody delabeling

Imaged SHIELD tissue was first equilibrated with the clearing buffer (200mM SDS, pH 9.5) at 37°C overnight. Afterwards the sample was moved to a separate falcon tube with 50 mL of clearing buffer that was preheated to 80°C and kept on a heated shaker maintained at 80°C for 1 h. Afterwards, the solution was exchanged with fresh clearing buffer at RT and the sample was incubated on an orbital shaker at 37°C overnight. The sample was washed using PBS with multiple solution exchanges for one day to thoroughly wash out SDS.

### Passive immunohistochemistry

Immunohistochemistry was performed on 100 μm- or 200 μm-thick mouse or marmoset brain tissue sections. Staining was performed on 24 or 48 well plates with primary antibodies (per recommended dilution from each vendors) and with dye-conjugated Fc-specific Fab fragments (3:1 molar ratio between Fab fragments and the primary antibody, Jackson Immunoresearch) for 1 day at RT in PBS with 0.1% Triton-X100. Similar protocols were used to characterize antibody binding performance in several different buffers: PBS with 0.1% NaDC, PBS with 1% NaDC, eFLASH initial buffer (240mM Tris, 160mM CAPS, 20% w/v D-sorbitol, 0.9% w/v NaDC, pH 9.6), and eFLASH terminal buffer (buffer retrieved from the eFLASH staining device after 24 h, pH 7.4).

### eFLASH protocol

Volumetric immunolabeling with eFLASH was carried out with a device described in Kim et al.^12^ Experiments were carried out with two buffers. The main buffer (240mM Tris, 160mM CAPS, 20% (w/v) D-sorbitol, 0.2% (w/v) NaDC) is a circulation solution that allows conduction of electricity. The sample buffer (240mM Tris, 160mM CAPS, 20% (w/v) D-sorbitol, 0.9% (w/v) NaDC) is used to fill the the sample cup along with the tissue and antibodies. 300 mL of a booster buffer (20% w/v D-sorbitol, 60mM Boric Acid) was added to the main buffer at 20 h after the start of the experiment to achieve the desired pH in the sample cup at 24 h.

300-500 mL of the main buffer was loaded into the staining device and 2-5 mL of the sample buffer was loaded into the sample cup. The tissue sample was placed in a nylon mesh then placed into the sample cup. Primary antibodies (antibody information and optimized quantity for each target, Supplementary Table 1) and secondary antibodies were added to the sample cup. Dye-conjugated Fc-specific Fab fragments were used for all experiments (2:1 molar ratio to the primary antibody, Jackson immunoresearch). The machine was operated for 24 h at 90 V with maximum current limited to 500 mA. Temperature control was set to maintain 25°C. Sample cup stir bar rotation was set to 850 rpm and sample cup rotation speed was set to 0.01 rpm.

### Dye conjugation of secondary antibodies

For the far-red channel, secondary antibodies conjugated with SeTau647 were used for most labeling experiments as they provide superior photo-stability when compared to commercially available dyes^62^. SeTau-647-NHS was purchased from SETA BioMedicals and 10 μl 10mM aliquots were prepared using DMSO (anhydrous, ZerO2®, ≥99.9%, Sigma). SeTau-647-NHS were reacted with non-conjugated Fc-specific Fab fragments at 10:1 ratio (Jackson Immunoresearch) for 1 h at RT. Afterwards, the solution was purified using Zeba Spin Desalting Columns (7k MWCO, ThermoFisher Scientific) 2 to 3 times until the desalting column ran clean. The concentration of the resulting solution was measured using DC™ Protein Assay (Bio-Rad) before use.

### Refractive index matching

Optical clearing of delipidated samples was achieved using Protos-based immersion medium^13^. For samples thicker than 1 mm, optical clearing was done in a step-wise manner. Labeled samples were first incubated in half-step solution (50/50 mix of 2X PBS and Protos-based immersion medium) at 37°C overnight. Afterwards, the samples were moved to the pure immersion medium and incubated at 37°C overnight.

### Fixation of labeled samples

For antibodies that are not stable in Protos-based immersion medium, the eFLASH-labeled samples were fixed with 4% (w/v) PFA to prevent dissociation of bound antibodies. eFLASH-labeled samples in the terminal labeling buffer were washed in 1X PBS with 0.02% (w/v) sodium azide at RT for at least 6 h to wash out Tris in the sample. Samples were then moved to freshly prepared 4% (w/v) PFA solution in 1X PBS, and placed on an orbital shaker at RT overnight. Samples were then washed with 1X PBS with 0.02% (w/v) sodium azide at RT with multiple solution exchanges for at least 6 h.

### Light-sheet imaging and post-processing

Rapid volumetric imaging was performed with an axially swept light-sheet microscope (SmartSPIM, Lifecanvas Technologies, MA) equipped with three lasers (488nm, 561nm, 642nm). The scanning was fine-tuned for each sample by finely adjusting the position of the illumination objectives to ensure optimal optical sectioning. Focus compensation was programmed as a function of depth for each laser line to account for slight focal variations through imaging depth. All light-sheet imaging was done with either of the following objective lenses: 3.6x objective (custom Lifecanvas design, 0.2NA 12mm WD lateral resolution 1.8um in XY), 10x objective (Olympus XLPLN10XSVMP, 0.6NA, 8mm WD, lateral resolution 0.66um in XY). Acquired data was post-processed with algorithms described in Swaney et al.^18^. A complete table of imaging modalities and conditions for every data included in this paper can be found in Supplementary table 2.

### Cell detection

Detection of cells is accomplished by blob detection, followed by dimensionality reduction and classification. Blobs are detected by computing the difference of Gaussians followed by identification of voxels that are the maximum of their neighbors within a chosen radius. 31×31 pixel patches are then extracted in the X/Y, X/Z and Y/Z planes. The rasters of these patches are concatenated and the three resulting 961-element vectors are concatenated to create a 2883-feature vector. All patches of putative cell centers within the volume are collected and PCA is performed to reduce the dimensionality of the vector to 48 components. Each of these components are composed of 2883 elements which are multiplied with the 2883-feature vector per patch to produce 48 numerical features. The vector of each component can be visualized as three 31×31 planes (see Supplementary Figure 3) to allow interpretation of the magnitude of the component. The 48 numerical features are then used to train a random forest classifier using iterative user-supervised training. Finally, the classifier is applied to all patches in the volume to classify each local maximum as a positive cell detection or negative artifact detection.

### Atlas Alignment

Atlas alignments of mouse brain hemispheres labeled with eFLASH to the Allen brain reference atlas, CCF V3^63^, were carried out using the hybrid automated atlas alignment method described in Swaney et al^18^, which combines Elastix^17^ and manual refinement tools to improve alignment accuracy.

### Brain region segmentation

Detected cell coordinates were transformed from the original coordinate space to the reference coordinate after atlas alignment. The alignment was used to construct a three-dimensional radial basis function using thin-plate spines to map points in the original coordinate space to the reference coordinate space. The point locations in the reference space were then matched against the Allen Brain Mouse Atlas reference^64^ segmentation to yield counts per brain region. These counts were then used to color the regions in the Allen Brain Mouse Atlas coronal SVG image files. Calculations and visualizations were done using the Nuggt python package^18^.

### Manual image analysis

Imaris (Bitplane, Switzerland) was used for soma segmentation, analysis, and neurite tracing in figure 5g-m. Dendrite polarity of NPY+ cells were assessed manually^65^. Fluorescence quantification was done using ImageJ.

## Code availability

The custom code used in this study is available from the corresponding author upon reasonable request.

## Data availability

The data supporting the findings of this study are available from the corresponding author upon reasonable request.

