## Supplementary figures and images for "Ultrafast immunostaining of organ-scale tissues for scalable proteomic phenotyping"

### Supplementary Figure 1

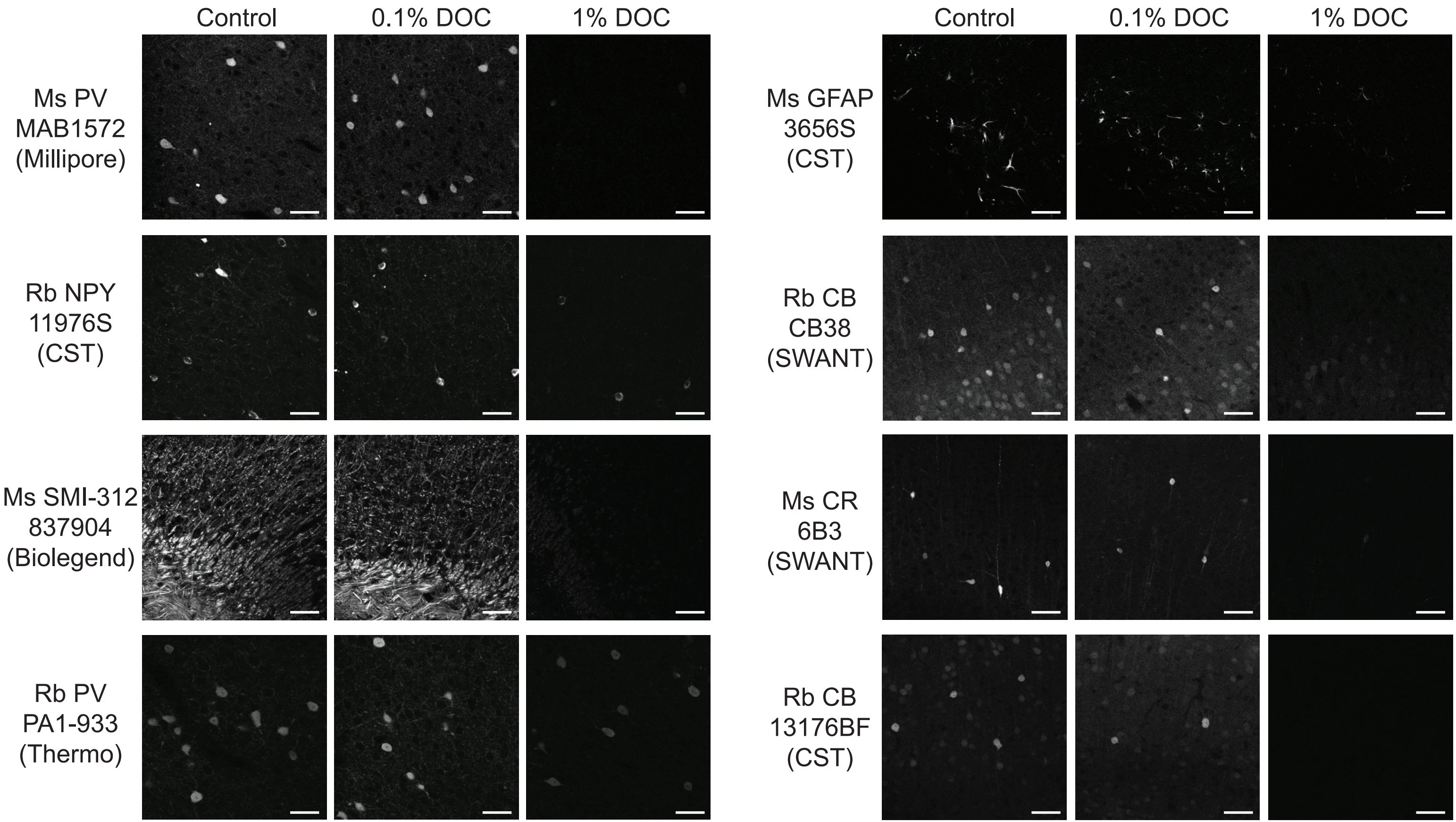

### Supplementary Figure 2

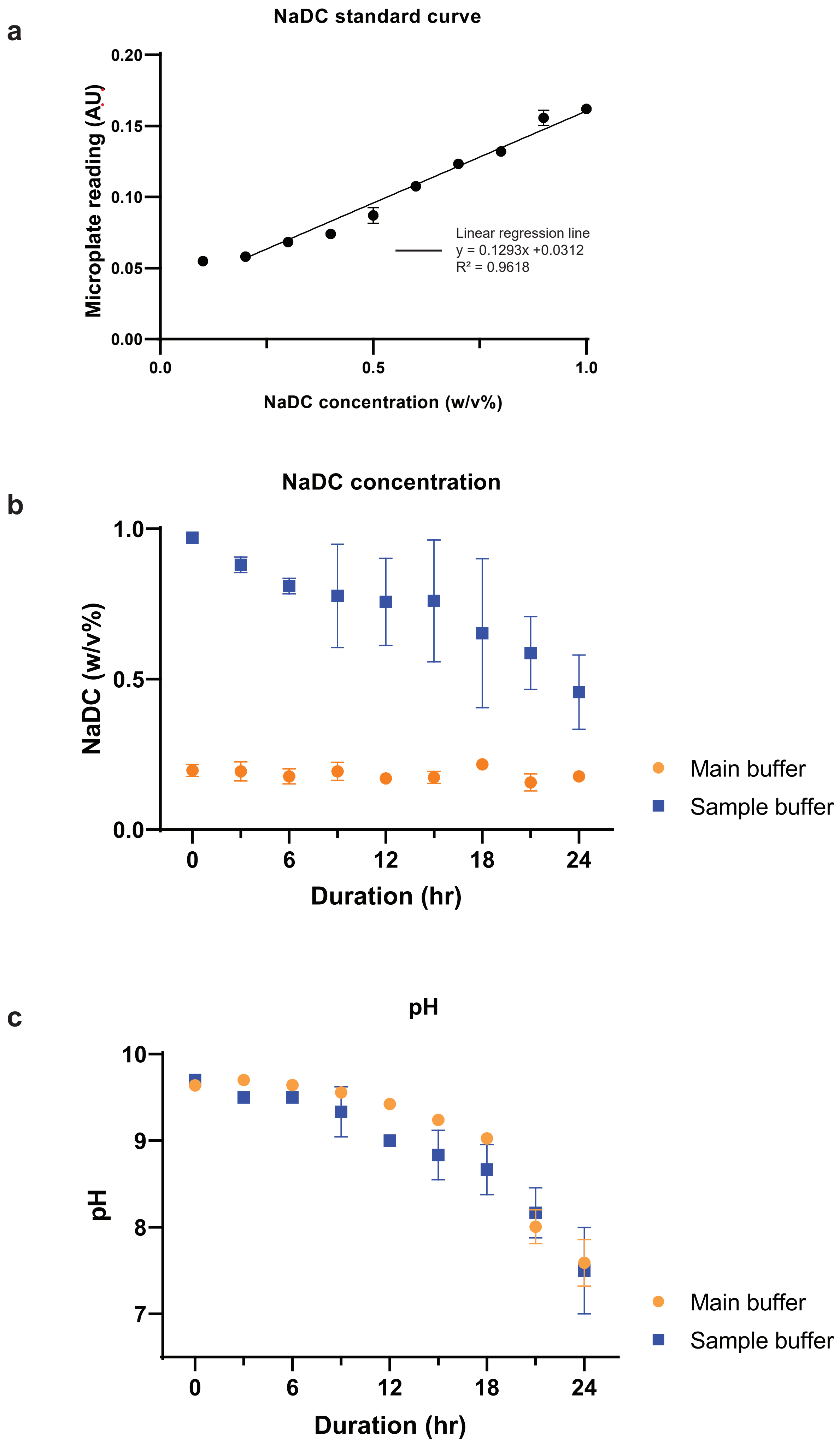

### Supplementary Figure 3

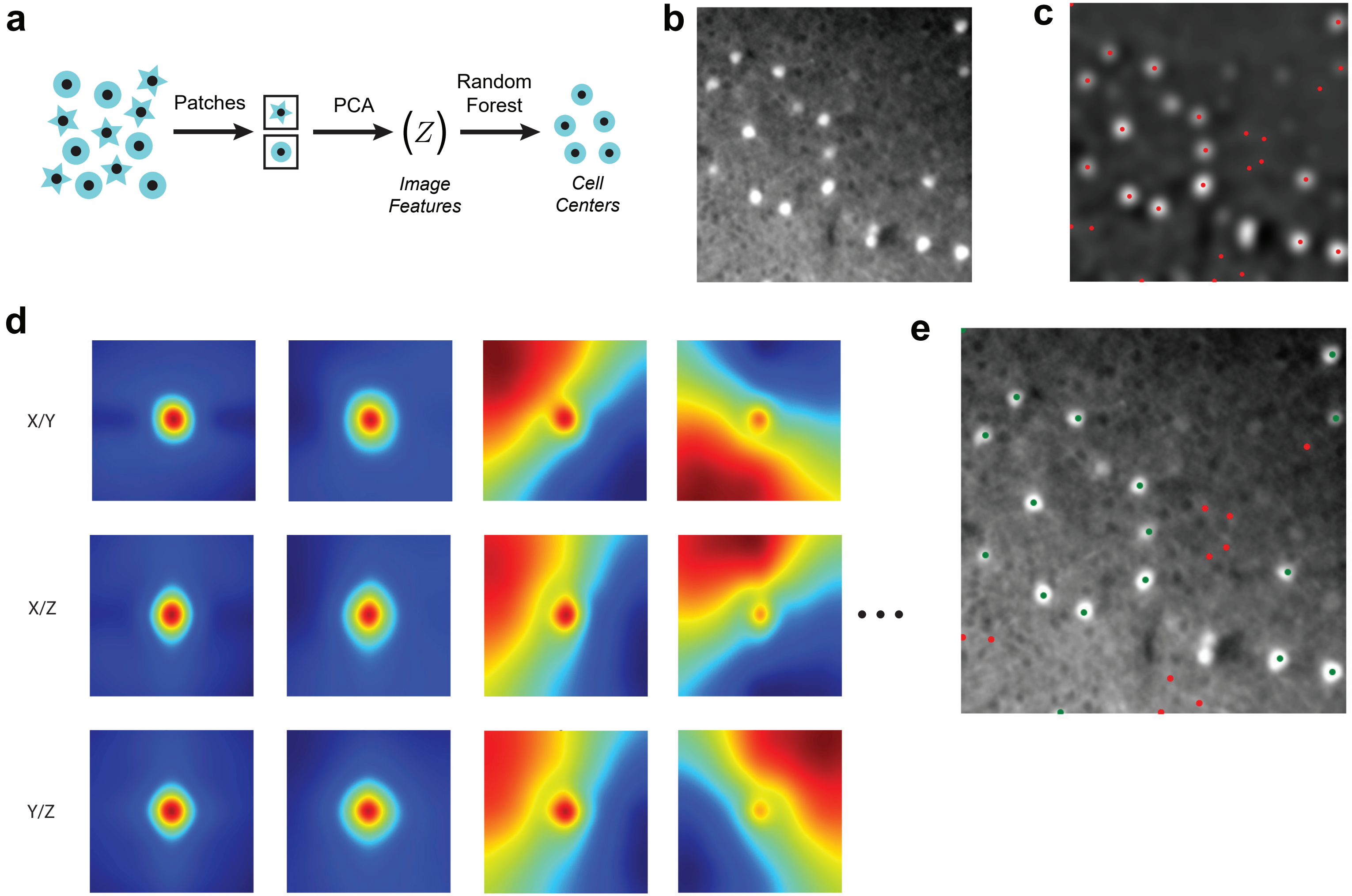

### Supplementary Figure 4

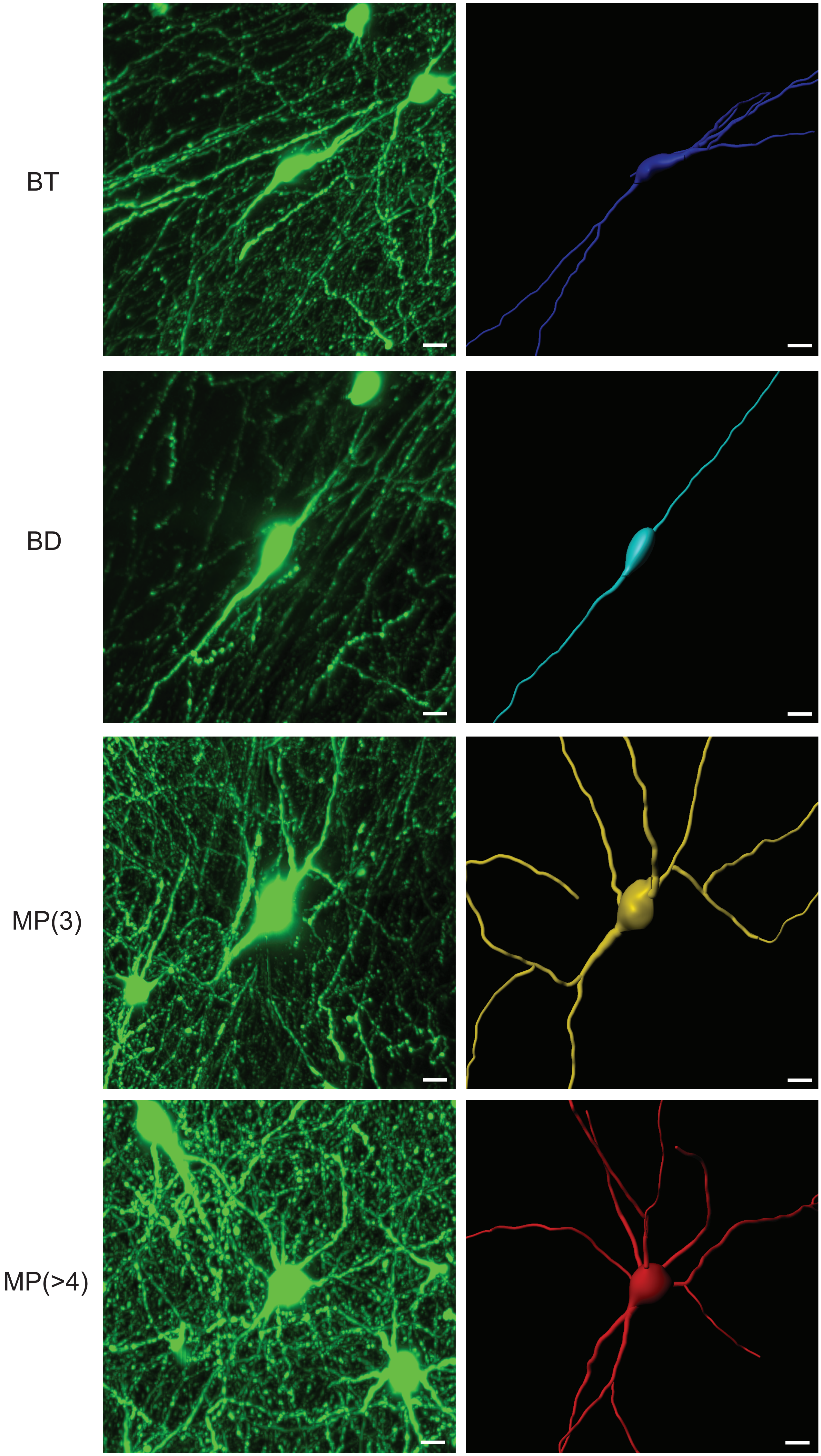
