## Supplementary Material Legends for "Ultrafast immunostaining of organ-scale tissues for scalable proteomic phenotyping"

**Supplementary Figure 1**. Antibody signals in the presence of sodium deoxycholate (NaDC). Diverse antibody signals were acquired under the control buffer (PBS with 0.1% Triton-X100) and 0.1% or 1% NaDC in PBS. All images of each antibody were acquired under the same condition.

**Supplementary Figure 2**. Measurement of NaDC concentration and pH of eFLASH buffers. (a) A standard curve for determining the concentration of NaDC in eFLASH bufferscollected at various time points during the experiment. Details can be found in the methods section. N = 3 replicate measurements. (b-c) NaDC concentrations (b) and pH (c) measurement for the two buffers used for eFLASH experiments (main and sample buffers) at 3 hour intervals. N = 3 independent experiments. Mean ± s.e.m.

**Supplementary Figure 3**. Cell detection algorithm. (a) Schematic of the algorithm. (b) Image of mouse brain stained with PV. (c) The difference of Gaussians is computed and local maxima are detected in 3 dimensions (red dots). (d) 31x31 patches in the X/Y, X/Z and Y/Z planes, centered at each local maxima are collected and PCA is performed to reduce the image to 48 components. The first and second components measure brightness at two different diameter scales and the third and fourth components measure intensity anisotropy in the X and Y directions. (e) A random-forest classifier is trained using the components as features and each maximum position is classified as a true (green dot) or false (red dot) detection.

**Supplementary Figure 4**. Representative NPY+ neurons with different dendrite polarities. Projection images of NPY+ neurons (left) and their reconstruction (right) were presented. BT, bitufted cell; BD, bidirectional cell; MP3, multipolar cell with three principal dendrites; MP (>4), multipolar cell with four or more numbers of principal dendrites. Scale bar = 20 µm.

**Supplementary Table 1**. Molecular probes used for eFLASH experiments. Optimized primary antibody amounts and molar ratios between primary antibody and secondary antibodies for each target are listed. Fc-specific Fab fragments from Jackson Immunoresearch was used for all eFLASH experiments. For antibodies with unknown concentrations, appropriate amount of secondary antibody was calculated based on empirically obtained data.

**Supplementary Table 2**. Sample preparation, imaging, and image processing conditions for all microscopy data presented in this paper.

**Supplementary Video 1.** Comparison between eFLASH-(left) and SE-(right) stained mouse brain hemispheres. Same amounts of antibodies were used for both conditions.

**Supplementary Video 2.** An eFLASH-stained mouse brain hemisphere with antibodies (anti-Parvalbumin and anti-SMI312) and a nuclei marker (syto16).

**Supplementary Video 3.** An eFLASH-stained marmoset brain block. A marmoset brain block (5 mm x 5 mm x 8 mm) containing the primary visual cortex was stained with anti-Neuropeptide-Y antibody.

**Supplementary Video 4.** Uniform volumetric immunolabeling of a mouse brain hemisphere with anti-Tyrosine Hydroxylase (TH), anti-Choline Acetyltransferase (ChAT), and anti-NeuN antibodies.

**Supplementary Video 5.** Uniform volumetric immunolabeling of a mouse brain hemisphere using eFLASH with an anti-cFos antibody. An adult mouse brain subjected to contextual fear conditioning was used.

**Supplementary Video 6.** Brain-wide detection of PV+ cells and its density heat map. An adult mouse brain hemisphere was labeled with anti-PV antibody with eFLASH. PV+ cells were automatically detected and the brain image was region-segmented to generate brain-wide a PV+ cell map.

**Supplementary Video 7.** PV-cre::DIO-tdTomato hemisphere eFLASH-stained with anti-PV antibody.

**Supplementary Video 8.** A $ChAT^{BAC}$-eGFP mouse brain hemisphere that was eFLASH-stained with anti-ChAT antibody.

**Supplementary Video 9.** Antibody labeling image (left) and PV+ cell density map of a marmoset visual cortical block (right). A marmoset brain block (5mm x 5mm x 8mm) containing visual cortical areas was stained with anti-PV antibody.

**Supplementary Video 10.** Dendrite polarity of NPY+ cells in marmoset V1
